# Inferred microbial networks lack replicability: consequences for next-generation biomonitoring

**DOI:** 10.1101/642199

**Authors:** Charlie Pauvert, Jessica Vallance, Laurent Delière, David A. Bohan, Marc Buée, Corinne Vacher

## Abstract

Plant-associated microbial interaction networks protect plants against disease. There is, therefore, a need to monitor in real time their responses to environmental changes to predict disease risk and adjust crop protection strategies. Next-Generation Biomonitoring (NGB) proposes to reconstruct automatically these networks from metabarcoding data, to complement ecological community properties commonly used for ecosystem health assessment. This study aimed to evaluate the benefits and shortcomings of community-level and network-level properties for biomonitoring. We specifically investigated whether microbial networks inferred from metabarcoding data show robust responses to agricultural practices, using the grapevine microbiota as a study system. Our results demonstrate a strong footprint of the agricultural practice on the metabarcoding data, when analyzed at the community level. The richness, diversity and evenness of fungal communities were significantly higher in organic than conventional plots. The cropping system also affected the composition of grapevine foliar fungal communities significantly. Contrary to our expectations, microbial networks were less sensitive to changes in agricultural practices than microbial communities, confirming that NGB should not only consider network-level properties but also community-level properties. Moreover, we found that microbial networks lacked replicability within a cropping system but that consensus networks, built from several network replicates, could generate relevant hypotheses of microbial interactions. As things stand, community-level properties appear to be a more reliable and statistically powerful monitoring option than network-level properties. Future developments, especially in network inference methods, are likely to challenge our findings and help to improve the monitoring of the ecosystem services provided by the plant microbiota.

## Introduction

Interactions among organisms and with their abiotic environment regulate the ecological processes underlying ecosystem services (Mace, Norris, & Fitter, 2012). Ecological interactions among organisms (e.g. predation, mutualism, parasitism) at a single point in space and time are usually represented as a network, with the organisms as nodes and the interactions as links (Pocock, Evans, & Memmott, 2012). Current challenges focus on understanding how and why these networks vary in space and time (Pellissier et al., 2018; Pilosof, Porter, Pascual, & Kéfi, 2017), and which network properties should be conserved or enhanced to sustain ecosystem services (Montoya, Rogers, & Memmott, 2012; Raimundo, Guimarães, & Evans, 2018; Tylianakis, Laliberté, Nielsen, & Bascompte, 2010). Next-Generation Biomonitoring (NGB) proposes to reconstruct ecological networks on top of community structures from Next-Generation Sequencing (NGS) data, and to analyze network and community variations in space and time for detecting and explaining changes in ecosystem functions and services (Baird & Hajibabaei, 2012; Bohan et al., 2017; Derocles et al., 2018).

NGB requires the reconstruction of replicated networks of ecological interactions as well as the development of statistical tools for comparison and analysis. Theoretical frameworks have been developed for the comparison of ecological networks between contrasting environmental conditions or along environmental gradients (Delmas et al., 2019; Pellissier et al., 2018; Poisot, Canard, Mouillot, Mouquet, & Gravel, 2012; Tylianakis & Morris, 2017). By analogy with the α- and β-diversity of ecological communities, these frameworks define α- and β-properties for ecological networks as whole-network metrics (e.g. connectance) and dissimilarities between pairs of networks, respectively (Pellissier et al., 2018). These metrics can be used to assess the impact of global environmental changes on the identity and abundance of the species forming ecological communities and on the type and strength of their interactions (Pecl et al., 2017; Scheffers et al., 2016). They can for instance be used to evaluate the impact of agricultural practices, which are a key driver of global change (Tilman, Cassman, Matson, Naylor, & Polasky, 2002), on species diversity (Tuck et al., 2014) and on pest and disease regulation services supported by species interactions (Ma et al., 2019; Macfadyen et al., 2009; Tylianakis, Tscharntke, & Lewis, 2007). Organic farming provides indeed lower levels of pests (Muneret et al., 2018) but others outcomes of such agriculture remains uncertain (Seufert & Ramankutty, 2017). Network metrics (Pellissier et al., 2018) could provide further insights into the footprint of agricultural practices, like organic farming, on the functioning of ecological communities and associated ecosystem services.

Networks of interactions among microorganisms are a natural target for NGB, because NGS techniques are the rule for studying microbial communities (Bálint et al., 2016), and because microbial networks are crucial to human life and well-being (Gilbert & Neufeld, 2014). Network ecology, which originates from the study of trophic links between macroorganisms (Ings et al., 2009), initially ignored interactions with and among smaller organisms (Lafferty, Dobson, & Kuris, 2006) despite evidence of the contribution of microbial interactions to biogeochemical cycles (Falkowski, Fenchel, & Delong, 2008), plant diversity and productivity (van der Heijden, Bardgett, & Straalen, 2008) and disease regulation (Berendsen, Pieterse, & Bakker, 2012). It is now recognized that microbial networks are particularly important for the health of animals and plants (i.e. holobionts; Zilber-Rosenberg & Rosenberg, 2008) because resistance to pathogens is mediated by direct antagonistic interactions between the residential microbiota and the pathogen species (i.e. the barrier effect; Arnold et al., 2003; Kamada, Chen, Inohara, & Núñez, 2013; Kemen, 2014; Koch & Schmid-Hempel, 2011; Laur et al., 2018) and by indirect interactions due to the activation of the host immune system by the residential microbiota (i.e. the priming effect; Hacquard, Spaepen, Garrido-Oter, & Schulze-Lefert, 2017; Kamada et al., 2013; Perazzolli et al., 2012; Ritpitakphong et al., 2016; Vogel, Bodenhausen, Gruissem, & Vorholt, 2016). The subset of a host-associated microbial network consisting of a pathogen and its interacting partners has been termed the pathobiome (Brader et al., 2017; Vayssier-Taussat et al., 2014). NGB of plant and animal health will require elucidation of the microbial interactions forming pathobiomes (Durán et al., 2018), identification of the intrinsic network properties that hinders invasion by pathogens (Agler et al., 2016; Murall et al., 2017; Poudel et al., 2016) and assessment of the changes in these properties (Creamer et al., 2016; Karimi et al., 2017; Morriën et al., 2017).

The success of NGB approaches will depend on our ability to automatically reconstruct networks that are similar to real ecological networks (Bohan, Caron-Lormier, Muggleton, Raybould, & Tamaddoni-Nezhad, 2011). Classically, ecological networks have been built on the basis of observations of ecological interactions. Metabarcoding approaches are now providing additional information (Evans, Kitson, Lunt, Straw, & Pocock, 2016), and making it possible to generate hypotheses about cryptic interactions such as those of microorganisms (Berry & Widder, 2014a; Faust & Raes, 2012). Hypotheses for microbial interactions can be formulated on the basis of co-abundance data derived from environmental DNA metabarcoding data (i.e. the sample × taxa matrix) via statistical or machine-learning approaches (Vacher et al., 2016). The representation of all positive and negative statistical associations between abundances for microbial taxa typically gives rise to a network resembling a hairball (Röttjers & Faust, 2018) and the challenge is to go beyond networks of this type, by removing spurious associations not due to microbial interactions and linking the remaining positive and negative associations to possible interaction mechanisms (Derocles et al., 2018). The first source of spurious associations is the compositional nature of metabarcoding data. In a metabarcoding dataset, the total number of sequences per sample is arbitrary, imposed by the sequencer. Sequence counts contain only relative abundance information for species. Comparisons that do not take this feature into account can result in the identification of artifactual associations (Gloor, Macklaim, Pawlowsky-Glahn, & Egozcue, 2017). Early methods of microbial network inference, such as SparCC (Friedman & Alm, 2012) attempted to overcome this bias using log ratios of counts. A second source of spurious associations is the joint response of microbial taxa to abiotic or biotic factors, creating indirect associations that reflect taxon-specific environmental requirements (Armitage & Jones, 2019; Röttjers & Faust, 2018). Early studies dealt with this issue by first using regression to eliminate environmental effects from the sequence counts and then inferring networks from the residuals (Biswas, Mcdonald, Lundberg, Dangl, & Jojic, 2016; Jakuschkin et al., 2016) and several methods integrating environmental covariates, such as HMSC (Ovaskainen et al., 2017), PLN (Chiquet, Mariadassou, & Robin, 2017, 2018), FlashWeave (Tackmann, Rodrigues, & Mering, 2018) and MAGMA (Cougoul, Bailly, & Wit, 2019), have since been developed. Finally, the taxonomic resolution of the nodes should be fine enough to discern the variation in ecological interactions between microbial strains (Röttjers & Faust, 2018). This has led to the generation of novel bioinformatics approaches that more fully exploit the resolution of molecular barcodes, such as DADA2 (Callahan et al., 2016). Despite these advances, however, the inference of microbial networks from metabarcoding data remains nascent (Layeghifard, Hwang, & Guttman, 2017), and inferred interactions should be interpreted with care ((Barner, Coblentz, Hacker, & Menge, 2018; Freilich, Wieters, Broitman, Marquet, & Navarrete, 2018; Röttjers & Faust, 2018; Weiss et al., 2016; Zurell, Pollock, & Thuiller, 2018), because few validation experiments have been conducted to date (e.g. Das, Ji, Kovatcheva-Datchary, Bäckhed, & Nielsen, 2018; Durán et al., 2018; Wang et al., 2017).

In this study, we investigated whether the impact of agricultural practices on microbial networks might be detected by combining current metabarcoding and network inference approaches. We first assessed the influence of conventional versus organic agriculture on plant-associated microbial communities using community-level properties and secondly using network-level properties. Furthermore, we checked whether microbial networks can generate robust hypotheses concerning microbial interactions. We inferred microbial networks from environmental DNA sampled from replicated agricultural plots and processed in DADA2 (Callahan et al., 2016) and SparCC (Friedman & Alm, 2012). Grapevine was the model plant species because effects of organic farming were detected on associated microbial communities (Castañeda, Miura, Sánchez, & Barbosa, 2018; Kernaghan, Mayerhofer, & Griffin, 2017; Pancher et al., 2012; Schmid, Moser, Muller, & Berg, 2011; Varanda et al., 2016) but remain to be assessed on networks. We focused on the fungal component of the foliar microbiota as it contains major pathogens (Armijo et al., 2016).

## Materials and methods

### Study site and sampling design

Samples were collected in 2015, from an experimental vineyard (Figure 1) located near Bordeaux (INRA, Villenave d’Ornon, France; 44°47’32.2”N 0°34’36.9”W). The experimental vineyard was planted in 2011 and was designed to compare three cropping systems: sustainable conventional agriculture (CONV), organic farming (ORGA) and pesticide-free farming (RESI) (Delière et al., 2014). The *Vitis vinifera* L. cultivar Merlot noir grafted onto a 3309 C rootstock was used in both the CONV and ORGA cropping systems. RESI used a disease-resistant cultivar and was not included in this study. The experiment had a randomized block design (Schielzeth & Nakagawa, 2013) consisting of three blocks, each composed of three plots, one for each of the cropping systems tested. Each plot covered an area of 2100 m^2^ and was composed of 20 rows of 68 vines each, with a 1.60 m between rows and 0.95 m between vines in a single row.

**Figure 1.**
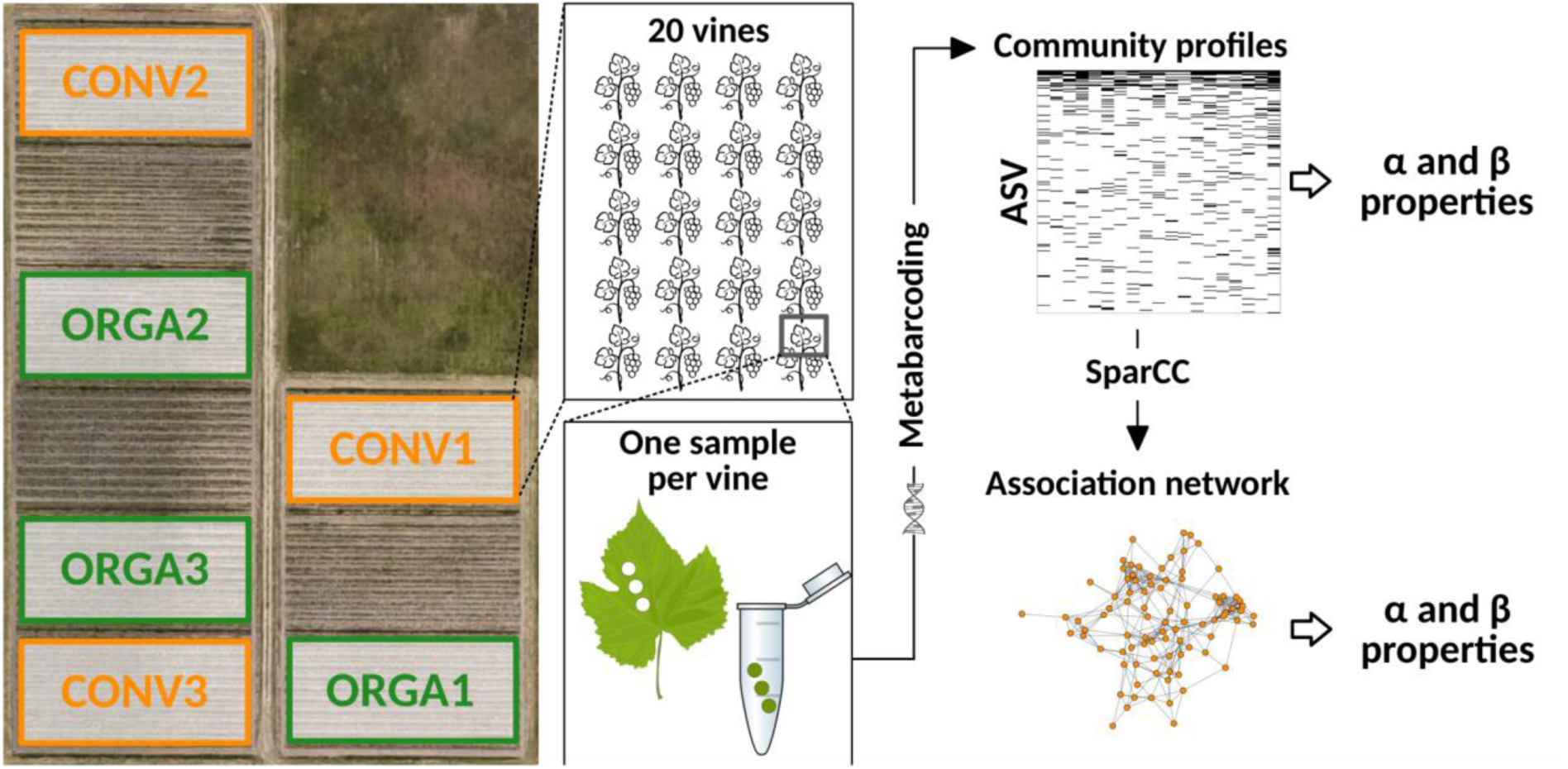
Experimental design. Foliar fungal communities were characterized in three conventional (CONV) and three organic (ORGA) vineyard plots by a metabarcoding approach. We analyzed 20 foliar samples per plot. For each plot, we thus obtained 20 community profiles (described in terms of amplicon sequence variants (ASV)) and one association network (inferred with the SparCC software developed by Friedman & Alm, 2012). More networks were then obtained by varying network reconstruction parameters (Figure 5). The effects of cropping system (CONV *versus* ORGA) on the grapevine foliar microbiota were assessed with both community and network α- and β-properties.

CONV plots were managed according to the general principles of integrated pest management (IPM), as listed in Appendix III of the 2009/128/EC Directive (European Commision, 2009). ORGA plots were managed according to European Council Regulation (EC) No 834/2007 (European Commision, 2007). ORGA plots were treated with copper and sulfur-based products, whereas additional phytosanitary products were allowed in CONV plots (Table S1). The cropping systems differed in terms of the types of pesticides applied and the timing of applications, but not in terms of doses (Table S1). All products and active ingredients were applied between the end of April and mid-August of 2015. Grapes were harvested on September 10, 2015. The disease incidence and severity at harvest were higher in CONV plots than in ORGA plots for both powdery mildew (caused by the fungal pathogen *Erysiphe necator*) and black rot (caused by the fungal pathogen *Guignardia bidwellii*). Downy mildew symptoms (caused by the oomycete pathogen *Plasmopara viticola*) did not differ significantly between the cropping systems (Table S2).

Grapevine leaves were collected a couple hours before grape harvest, from 20 vines per plot in the CONV and ORGA plots (Figure 1). We attempted to avoid edge effects by selecting the 20 vines from the center of each plot. The third leaf above the grapes was collected from each vine, placed in an individual bag and immediately transported to the laboratory. In total, 120 leaves, corresponding to 1 leaf × 20 vines × 3 plots × 2 cropping systems, were collected. Leaves were processed on the day of collection, with sterilized tools in the sterile field of a MICROBIO electric burner (MSEI, France). Three contiguous discs of 6 mm diameter were cut from the center of each leaf, approximately 2 cm from the midrib. They were placed in the well of a sterile DNA extraction plate. The leaf disks were then freeze-dried overnight (Alpha 1-4 DA Plus, Bioblock Scientific).

### DNA extraction and sequencing

Leaf disks (Figure 1) were ground with a single-glass ball mill (TissueLyser II, Qiagen) and DNA was then extracted with a CTAB chloroform/isoamyl alcohol (24:1) protocol. A dozen “empty” wells (*i.e.* containing nothing but extraction reagents) were included on each plate as negative control samples for DNA extraction. Three of these negative control samples were randomly selected and pooled before sequencing. Three replicates of a fungal mock community, each consisting of an equimolar pool of DNA from 189 pure fungal strains, were also included as positive control samples (Pauvert et al., 2019).

The nuclear ribosomal internal transcribed spacer (ITS) region, which is considered to be the universal barcode region for fungi (Schoch et al., 2012), was then amplified with the ITS1F (5’-CTTGGTCATTTAGAGGAAGTAA-3’, Gardes & Bruns, 1993) and ITS2 (5’-GCTGCGTTCTTCATCGATGC-3’, White, Bruns, Lee, & Taylor, 1990) primer pair, which targets the ITS1 region. PCR was performed in an Eppendorf thermocycler (Eppendorf), with a reaction mixture (25 µl final volume) consisting of 0.04 U *Taq* polymerase (SilverStar DNA polymerase, Eurogentec), 1X buffer, 2 mM MgCl_2_, 200 µM of each dNTP, 0.2 µM of each primer, 1 ng.µl^-1^ bovine serum albumin (New England BioLabs) and 2 µl DNA template. A pseudo-nested PCR protocol was used, with the following cycling parameters: enzyme activation at 95°C for 2 min; 20 (1^st^ PCR with regular primers; Table S3) and then 15 (2^nd^ nested PCR with pre-tagged primers; Table S3) cycles of denaturation at 95°C for 30 s, 53°C for 30 s, 72°C for 45 s; and a final extension phase at 72°C for 10 min. “Empty” wells (i.e. containing nothing but PCR reagents) were included on each plate as a negative control for PCR. Three negative control samples were randomly selected and pooled before sequencing. In addition, the six PCR products corresponding to the 24^th^ leaf of the six plots under study (three CONV plots, three ORGA plots) were split in two, with each half of the sample sequenced independently to serve as technical replicates for sequencing.

We checked the quality of all the PCR products by electrophoresis in 2% agarose gels. PCR products were purified (CleanPCR, MokaScience), multiplex identifiers and sequencing adapters were added, and library sequencing on an Illumina MiSeq platform (v3 chemistry, 2×250 bp) and sequence demultiplexing (with exact index search) were performed at the Get-PlaGe sequencing facility (Toulouse, France).

### Bioinformatic analysis

Based on the mock community included in the sequencing run, we found that analyzing single forward (R1) sequences with DADA2 (Callahan et al., 2016) was a good option for fungal community characterization (Pauvert et al., 2019). Using DADA2 v1.6, we retained only R1 reads with less than one expected error (based on quality scores; Edgar & Flyvbjerg, 2015) that were longer than 100 bp, and we then inferred amplicon sequence variants (ASV) for each sample. Chimeric sequences were identified by the consensus method of the *removeBimeras* function. Taxonomic assignments were performed with RDP classifier (Wang et al., 2007), implemented in DADA2 and trained with the UNITE database v. 7.2 (UNITE Community 2017). Only ASVs assigned to a fungal phylum were retained. The ASV table was then filtered as described by Galan et al., (2016) with a script (https://gist.github.com/cpauvert/1ba6a97b01ea6cde4398a8d531fa62f9) that removed ASVs from all samples for which the number of sequences was below the cross-contamination threshold, defined as their maximum number in negative control samples. Finally, we checked the compositional similarity of the technical replicates (Figure S1), retaining the sample with the largest number of sequences for each replicate. The final ASV table contained 1116 ASVs, 112 samples and 4,760,068 high-quality sequences.

### Statistical analyses

Statistical analyses were performed with R software v3.4.1 (R Core Team, 2017), with the packages lme4 (Bates, Mächler, Bolker, & Walker, 2015), vegan (Oksanen et al., 2018), permute (G. L. Simpson, 2016), phyloseq (McMurdie & Holmes, 2013) including the DESeq2 extension (Love, Huber, & Anders, 2014), and igraph (Csardi & Nepusz, 2006). Data were manipulated and plots were created with reshape2, plyr and ggplot2 (Wickham, 2007, 2011, 2016), cowplot (Wilke, 2018), ggraph (Pedersen, 2018) and VennDiagram (Chen, 2018).

#### Effect of cropping system on community α-diversity

Generalized linear mixed models (GLMMs) were used to test the effect of cropping system on the richness, diversity and evenness of fungal communities. The models included the cropping system as a fixed treatment effect, the block and its interaction with the cropping system as random factors, and the sampling depth (defined as the total number of raw sequences per sample) as an offset (Bálint et al., 2015; McMurdie & Holmes, 2014). Community richness was defined as the number of ASVs per sample. We used a logarithmic link function to model these count data, assuming a negative binomial distribution to deal with overdispersion (Zuur, Ieno, Walker, Saveliev, & Smith, 2009). Community diversity was measured with the Inverse Simpson index (E. A. Simpson, 1949) and modeled with a Gaussian distribution and the logarithmic link function. Evenness was estimated with Pielou’s index (Pielou, 1966) and modeled with a Gaussian distribution and the logarithmic link function. The offset was transformed according to the link function. The significance of the fixed treatment effect was finally assessed with the Wald χ^2^ test (Bolker et al., 2009).

#### Effect of cropping system on community β-diversity

Permutational analyses of variance (PERMANOVAs; Anderson, 2001) were used to evaluate the effect of cropping system on compositional dissimilarities between fungal communities detected with the quantitative and binary versions of the Jaccard dissimilarity index (Chao, Chazdon, Colwell, & Shen, 2006; Jaccard, 1900). The models included cropping system, sampling depth (log-transformed) and their interaction as fixed effects. Permutations (*n* = 999) were constrained within blocks. ASVs differing in abundance between cropping systems were identified with DESeq2 (Love et al., 2014), by calculating the likelihood ratio between a full model including block and cropping system as fixed effects and a simplified model including only the block factor. The estimated fold-changes in abundance were considered significant if the *p*-value was below 0.05 after Benjamini and Hochberg adjustment.

#### Effect of cropping system on network α-properties

Fungal association networks were inferred at plot level (Figure 1) with the SparCC algorithm (Friedman & Alm, 2012) implemented in FastSpar (Watts, Ritchie, Inouye, & Holt, 2019) with default SparCC values. Ten networks per plot were constructed by varying the percentage *P* of ASVs included in the network (with *P* ranging from 10% to 100% of the most abundant ASVs in the plot). Networks had ASVs as nodes and a positive or negative link between ASVs in cases of significant associations between abundance. Six *α-*properties were calculated for all networks: number of links, network density, number of connected components, diameter of the largest component, mean node degree and proportion of negative links (Table S4). The effect of cropping system on these properties was investigated by performing Wilcoxon rank-sum tests for all values of *P*. The Benjamini-Hochberg procedure was used to correct *p*-values for multiple testing.

#### Effect of cropping system on network β-properties

The topological distance between networks was calculated for all pairs of networks, with the *D* index defined by Schieber et al. (2017) for binary networks. The dissimilarity of associations between networks, β_WN_ according to the framework described by Poisot et al. (2012), was then calculated for all pairs of networks with the binary Jaccard dissimilarity index. β_WN_ was then partitioned into two components (Poisot et al., 2012): the dissimilarity of associations between ASVs common to both networks (β_OS_) and the dissimilarity of associations due to species turnover (β_ST_). PERMANOVA was used to evaluate the effect of cropping system on the topological distance between networks (*D*) and the dissimilarity of associations between networks (β_WN_, β_ST_ and β_OS_). The models included cropping system, *P* and their interactions as fixed effects. The permutations (*n*=999) were constrained within blocks. Consensus networks were built to identify robust associations that could indicate biotic interactions between fungal strains.

## Results

The foliar fungal communities were dominated by Ascomycota in both ORGA (87.2% of sequences) and CONV (96.8%) plots. They were largely colonized by an ASV assigned to the *Aureobasidium* genus. More than half the sequences belonged to this ASV (Table 1), whatever the cropping system. The causal agent of grapevine powdery mildew, *Erysiphe necator*, was among the 10 most abundant fungal species. The proportion of sequences assigned to this pathogen species was higher in CONV than in ORGA plots (Table 1), consistent with the visual records of disease symptoms (Table S2).

**Table 1.** Most abundant amplicon sequence variants (ASVs) in grapevine foliar fungal communities according to cropping system. The relative abundances (RA, in %) and ranks of ASVs were calculated for all leaf samples (TOTAL; *n* = 112) and for samples collected from organic (ORGA; *n* = 55) and conventional plots (CONV; *n* = 57).

| ASV taxonomic assignment | TOTAL |  | ORGA |  | CONV |  |
| --- | --- | --- | --- | --- | --- | --- |
|  | Rank | RA | Rank | RA | Rank | RA |
| *Aureobasidium sp.* | 1 | 61.4 | 1 | 55.8 | 1 | 66.7 |
| *Cladosporium delicatulum* | 2 | 6.3 | 4 | 6.9 | 2 | 5.8 |
| *Filobasidium sp.* | 3 | 5.1 | 2 | 9.7 | 9 | 0.7 |
| *Alternaria sp.* | 4 | 4.4 | 5 | 3.9 | 4 | 5.0 |
| *Epicoccum nigrum* | 5 | 4.1 | 7 | 2.7 | 3 | 5.4 |
| *Cladosporium ramotenellum* | 6 | 3.5 | 3 | 7 | 46 | <0.1 |
| *Mycosphaerella tassiana* | 7 | 3.3 | 8 | 1.8 | 5 | 4.8 |
| *Didymella sp.* | 8 | 1.4 | 6 | 2.7 | 33 | 0.1 |
| *Erysiphe necator* | 9 | 1.1 | 38 | <0.1 | 6 | 2 |
| *Vishniacozyma victoriae* | 10 | 0.9 | 9 | 1.6 | 17 | 0.3 |

### Effect of cropping system on community α- and β-diversity

Fungal community richness, diversity and evenness were significantly higher in ORGA than CONV plots (Wald χ^2^ = 4.74, *p* = 0.029; Wald χ^2^ = 8.28, *p* = 0.004; Wald χ^2^ = 12.88, *p <* 0.001, respectively; Figure 2). The composition of foliar fungal communities differed significantly between cropping systems (Table 2), in terms of both the relative abundance (Figure 3A) and presence-absence of ASVs (Figure 3B). DESeq2 analysis revealed that four ASVs, including the fungal pathogen *Erysiphe necator*, were significantly more abundant in CONV plots, whereas 10 other ASVs, including several yeast species (from the genera *Vishniacozyma, Sporobolomyces* and *Filobasidium*), were significantly more abundant in ORGA plots (Figure 3C). Principal coordinate analysis (PCoA) with the Jaccard quantitative index (Figure 3A) revealed large differences in the relative abundances of ASVs between samples within a cropping system. The first axis of the PCoA accounted for 32.5% of the variance in community composition but did not discriminate between cropping systems. It was significantly correlated with the relative abundance of the dominant ASV (assigned to the *Aureobasidium* genus) (Spearman *ρ =* 0.96; *p* < 0.001).

**Table 2.** Effect of cropping system — conventional *versus* organic — on the β-diversity metrics of grapevine foliar fungal communities. Dissimilarities in community composition between samples were assessed with both the quantitative and binary Jaccard indices. The effects of sequencing depth (SD, log-transformed) and cropping system on composition dissimilarities between communities were evaluated in permutational analyses of variance (PERMANOVA). The number of permutations was set to 999 and permutations were constrained by block.

| Dissimilarity index | PERMANOVA |  |  |  |  |
| --- | --- | --- | --- | --- | --- |
| Quantitative<br>Jaccard | Variable | Df | F | R2 | Pr(>F) |
|  | Sequencing_Depth (SD) | 1 | 4.49 | 0.04 | <0.01 |
|  | Cropping_System (CS) | 1 | 9.42 | 0.08 | <0.01 |
| | SD × CS | 1 | 1.17 | 0.01 | 0.26 |
|  | Residuals | 108 |  | 0.854 |  |
|  | Total | 111 |  | 1 |  |
| Binary<br>Jaccard | Variable | Df | F | R2 | Pr(>F) |
|  | Sequencing_Depth (SD) | 1 | 1.05 | 0.01 | 0.29 |
|  | Cropping_System (CS) | 1 | 5.21 | 0.05 | <0.01 |
| | SD × CS | 1 | 1.03 | 0.01 | 0.41 |
|  | Residuals | 108 |  | 0.937 |  |
|  | Total | 111 |  | 1 |  |

**Figure 2.**
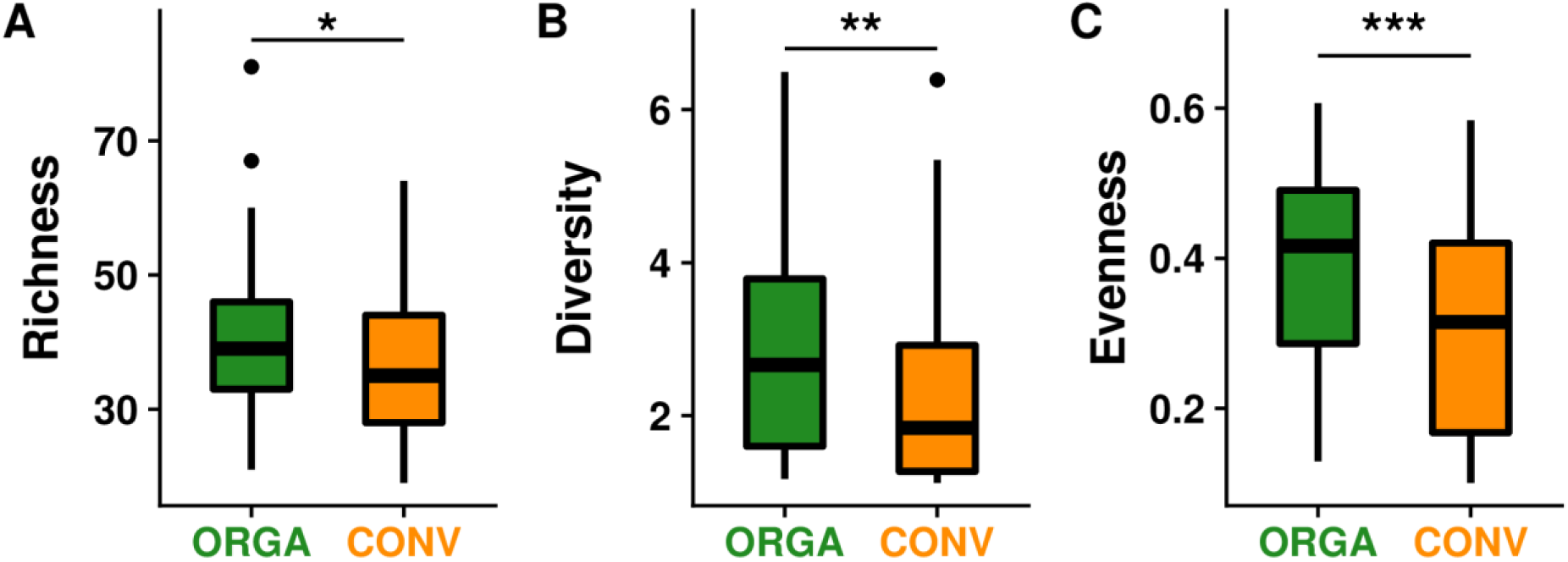
Effect of cropping system —conventional (CONV) versus organic (ORGA) — on the α-diversity metrics of grapevine foliar fungal communities. (A) Community richness, defined as the number of ASVs. (B) Community diversity, measured with the inverse Simpson index. (C) Community evenness, measured with Pielou’s index. Differences in α-diversity metrics between cropping systems were evaluated in Wald χ^2^ tests (* p<0.05; **p<0.01; ***p<0.001).

**Figure 3.**
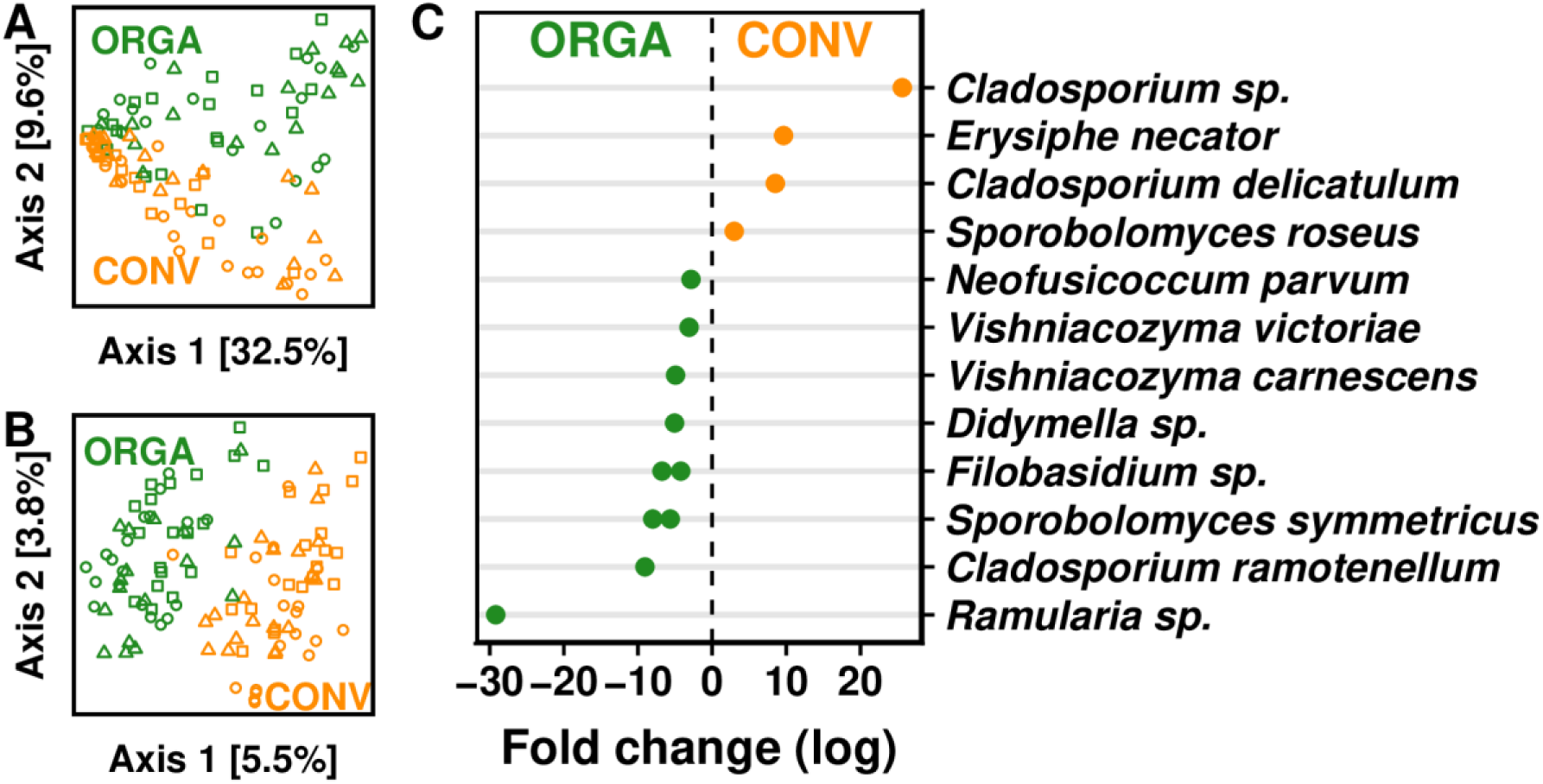
Effect of cropping system — conventional (CONV) *versus* organic (ORGA) — on the β-diversity metrics of grapevine foliar fungal communities. Principal coordinate analyses (PCoA) were used to represent dissimilarities in composition between samples, as assessed with the (A) quantitative and (B) binary Jaccard indices. The effect of cropping system on both β-diversity metrics was significant (Table 2). Green circles, squares and triangles correspond to samples collected in the ORGA1, ORGA2 and ORGA3 plots, respectively. Orange circles, squares and triangles correspond to the CONV1, CONV2 and CONV3 plots, respectively (Figure 1). (C) Log-transformed ratio of ASV relative abundance in CONV plots over that in ORGA plots, for 14 ASVs identified as differentially abundant between cropping systems by DESeq2 analysis followed by Benjamini-Hochberg adjustment (Love et al., 2014).

### Effect of cropping system on network α- and β-properties

In total, we obtained sixty fungal association networks, corresponding to the ten values of *P* (Figure 4A) for each of the six plots (Figure 1). None of the six network α-properties (Table S4) differed between cropping systems (Table S5), but all were significantly correlated with the percentage *P* of ASVs included in the network (Table S6). The total number of links and the mean node degree increased with *P*, whereas the number of connected components, network diameter, connectance and the proportion of negative links decreased (Figure 4B; Table S6). Similarly, the topological distance between networks did not differ between cropping systems, but was influenced by *P* (Table 3). By contrast, cropping system had a significant effect on the overall dissimilarity of associations (β_WN_) and the dissimilarity of associations between shared ASVs (β_OS_; Table 3 and Figure 5). However, fungal association networks varied considerably between plots within a cropping system (Figure 4A and Figure 6). When all ASVs were used for network construction (*P*=100%), only two associations were common to all three network replicates of the ORGA system (Figure 6A; Figure S2A) and only six were common to all three network replicates of the CONV system (Figure 6B; Figure S2B). The two associations replicated in the ORGA system were negative associations between the dominant ASV (assigned to the genus *Aureobasidium*) and *Cladosporium ramotenellum*, and between *Vishniacozyma victoriae* and *Neofusicoccum parvum* (Figure S2A). Five of the six associations replicated in the CONV system were positive, the remaining negative association being that between the dominant ASV (assigned to the genus *Aureobasidium*) and *Epicoccum nigrum* (Figure S2B). No association common to all six networks was identified (Figure 6C).

**Table 3.**
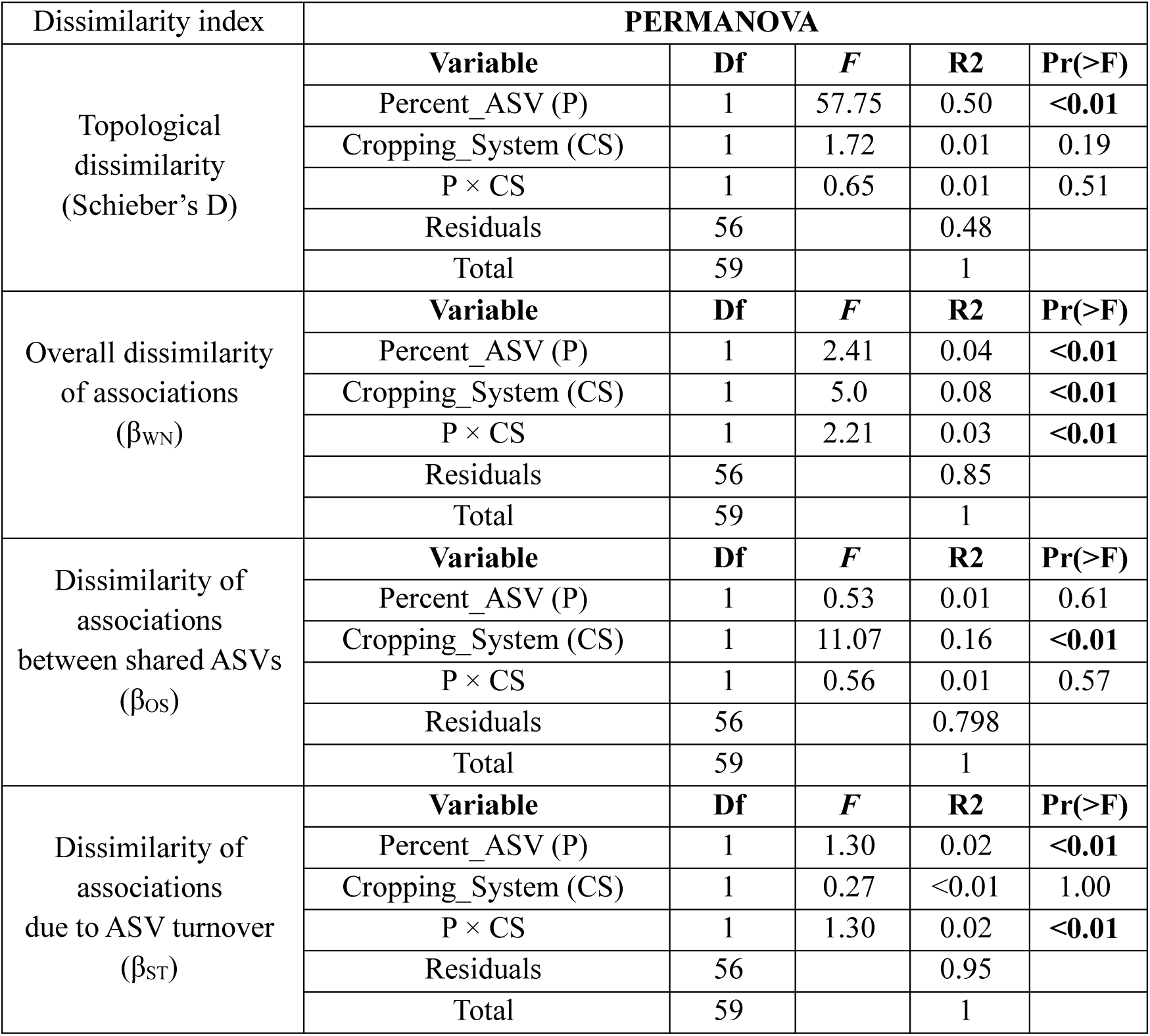
Effect of cropping system — conventional *versus* organic — on the β-properties of grapevine foliar fungal networks. The D index quantifies the topological dissimilarity between networks (Schieber et al., 2017) whereas the other three metrics (β_WN_, β_OS_ and β_ST_), which were calculated with the binary Jaccard index, quantify differences in associations between networks (Poisot et al., 2012). The effect of the percentage *P* of the most abundant ASVs used for network inference, and the effect of cropping system on the dissimilarities between networks were evaluated in permutational analyses of variance (PERMANOVA). The number of permutations was set to 999 and permutations were constrained by block.

**Figure 4.**
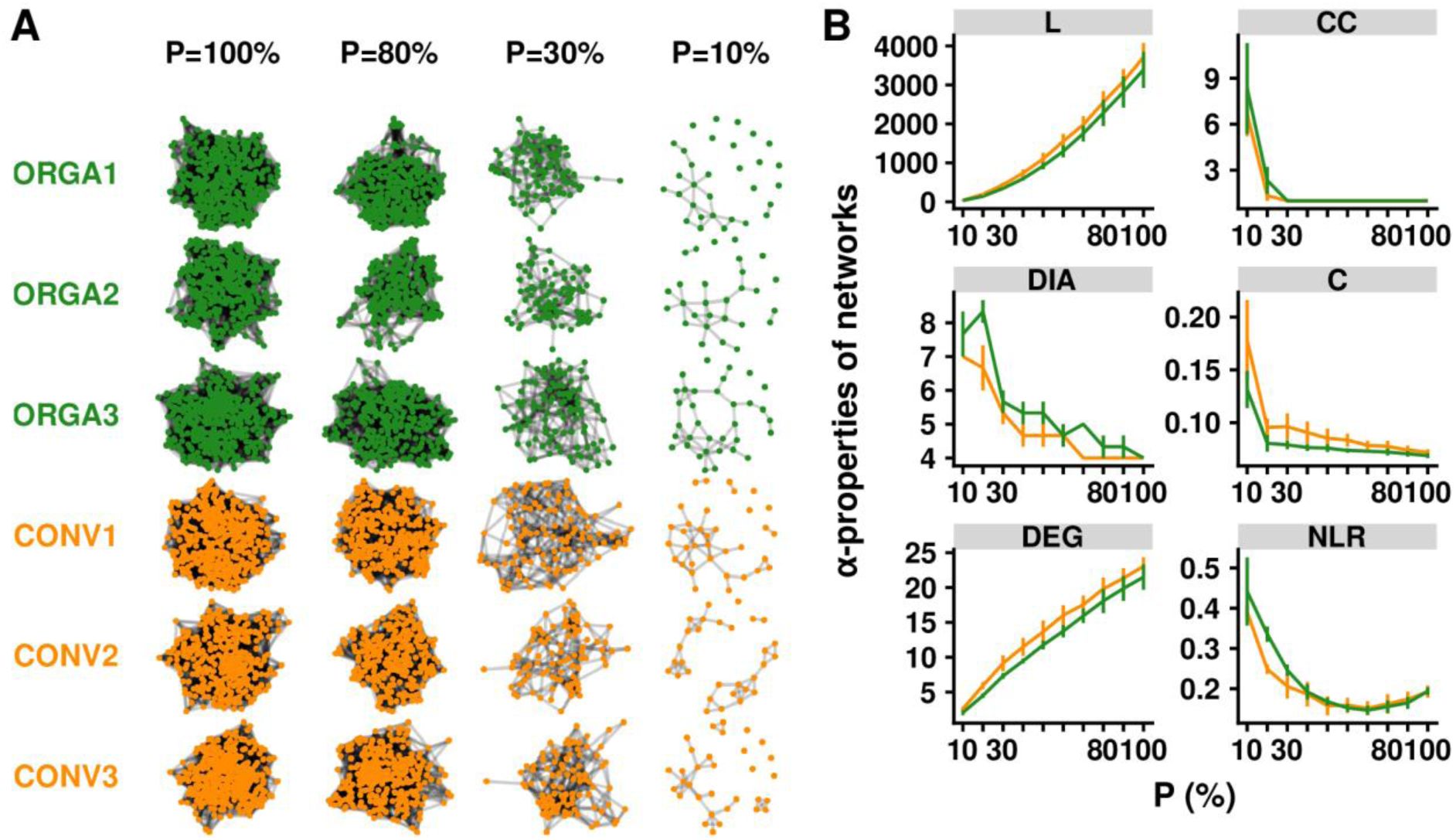
Effect of cropping system — conventional (CONV) *versus* organic (ORGA) — on the α-properties of grapevine foliar fungal networks. (A) Association networks inferred from fungal metabarcoding data with SparCC (Friedman & Alm, 2012). A total of 60 networks were inferred, corresponding to 2 cropping systems × 3 replicates (blocks) × 10 *P* values, with *P* the percentage of most abundant ASVs used for network inference. Only four values of *P* are shown on the figure. (B) Variations in network α-properties. The following properties (Table S4) were calculated for each network: the number of links (L) and connected components (CC), the network diameter (DIA) and connectance (C) and the mean degree (DEG) and negative link ratio (NLR). The percentage *P* of ASVs used for network reconstruction had a significant influence on all properties (Table S6), whereas cropping system did not (Table S5).

**Figure 5.**
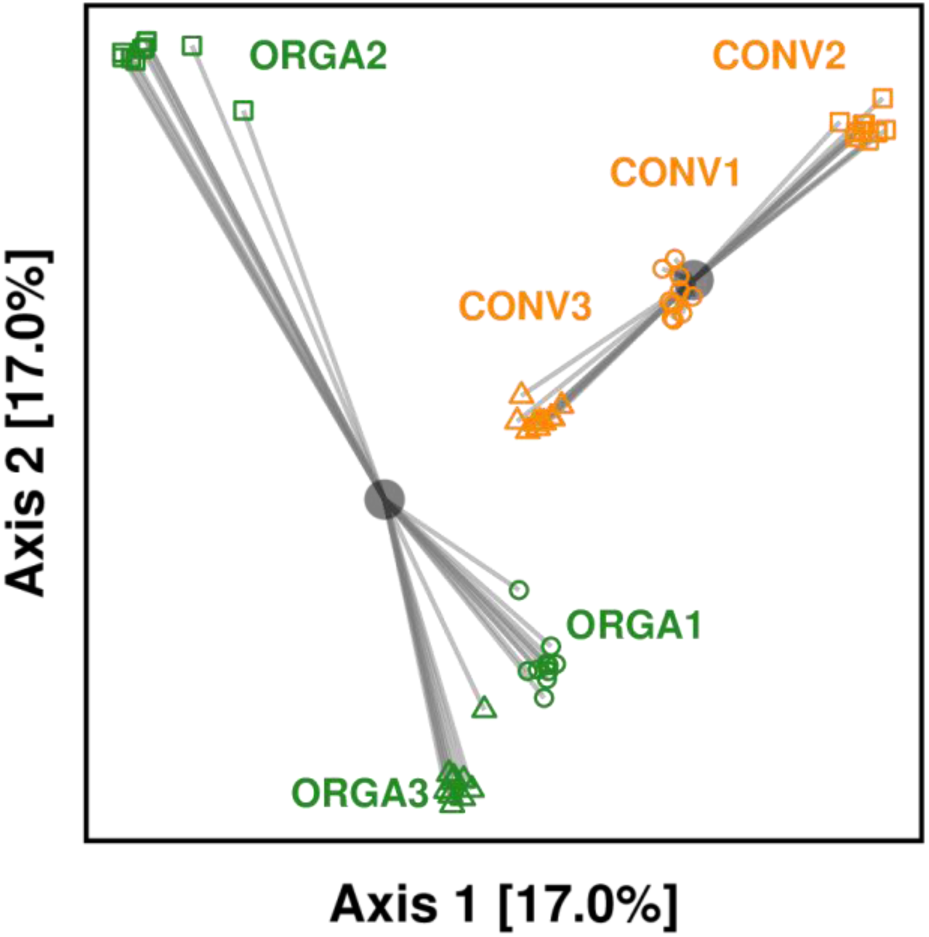
Effect of cropping system —conventional (CONV) *versus* organic (ORGA) — on the β-properties of grapevine foliar fungal networks. Principal coordinate analysis (PCoA) representing dissimilarities between networks, measured with the β_OS_ index (Poisot et al., 2012) calculated with the binary Jaccard index. β_OS_ measures the dissimilarity between two networks in terms of the presence-absence of associations between shared ASVs. The centroids for each cropping system are represented by gray circles. The effect of cropping system on β_OS_ was significant (Table 3).

**Figure 6.**
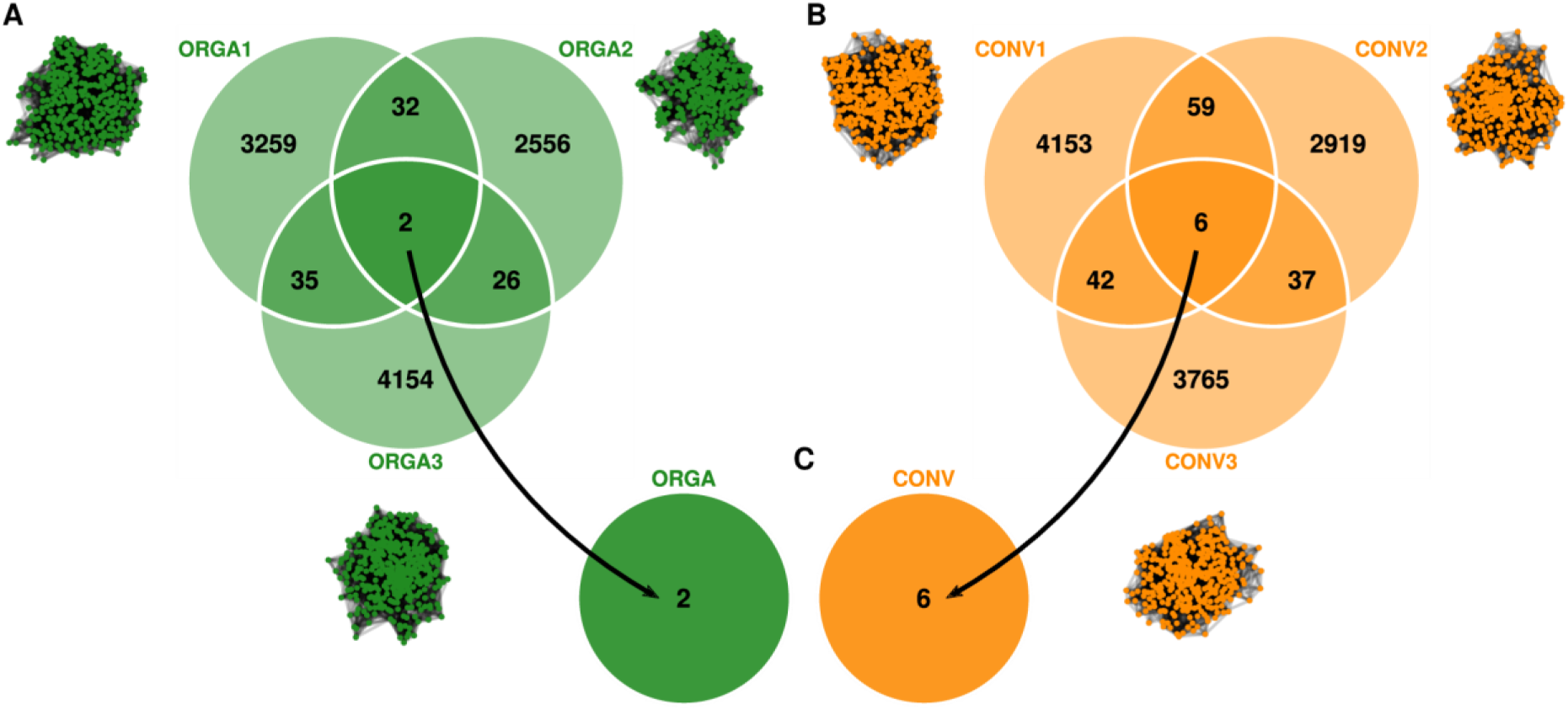
Venn diagrams showing the number of fungal associations common to network replicates. (A) Associations common to the three network replicates inferred for the organic cropping system (ORGA1, ORGA2, ORGA3) and (B) the three network replicates inferred for the conventional cropping system (CONV1, CONV2, CONV3), regardless of the sign of the association, in the situation in which all ASVs were used for network construction (*P*=100%). (C) Associations common to the six networks.

## Discussion

Plant-associated microbial interaction networks protect plants against disease (Hassani, Durán, & Hacquard, 2018; Kemen, 2014). Their responses to environmental changes, such as changes in agricultural practices, must be monitored in real time to better forecast disease risk. The concept of Next-Generation Biomonitoring (NGB) proposes that this could be done via the automatic reconstruction of ecological networks from metabarcoding data (Bohan et al., 2017). Therefore, we investigated whether automatically-reconstructed microbial networks have robust responses to agricultural practices (conventional *versus* organic agriculture). Using grapevine foliar fungal communities as study system, we found a strong footprint of the agricultural practice on the metabarcoding data, when analyzed at the community level. The richness, diversity and evenness of fungal communities were significantly higher in organic than conventional vineyards, consistent with the recent findings of Kernaghan et al. (2017) (but see Castañeda et al., 2018). The cropping system also significantly affected the composition of grapevine foliar fungal communities, as reported in previous studies (Castañeda et al., 2018; Kernaghan et al., 2017; Pancher et al., 2012; Schmid et al., 2011; Varanda et al., 2016). For instance, *Erysiphe necator*, the causal agent of grapevine powdery mildew, was significantly more abundant in conventional than in organic plots. The cause for such contrast in the pathogen abundance remains unknown given the absence of difference in dose and number of treatments. These results are, however, consistent with visual assessments of disease symptoms, indicating that, despite their numerous biases, metabarcoding data do contain some quantitative information useful for monitoring plant disease development (Jakuschkin et al., 2016; Makiola et al., 2018; Sapkota, Knorr, Jørgensen, O’Hanlon, & Nicolaisen, 2015). Several yeast strains, assigned to the genera *Vishniacozyma, Sporobolomyces* and *Filobasidium*, were significantly more abundant in organic plots. These yeast genera are frequently detected on leaf surfaces due to their tolerance of irradiation and they might influence plant growth by producing plant hormone-like metabolites (Kemler, Witfeld, Begerow, & Yurkov, 2017). In addition, *Vishniacozyma victoriae* (ex *Cryptococcus victoriae*) was reported as a biocontrol agent on postharvest diseases (Lutz, Lopes, Rodriguez, Sosa, & Sangorrín, 2013). Others yeasts possess wanted features of such agents like killer activities for some *Sporobolomyces* yeasts (Klassen, Schaffrath, Buzzini, & Ganter, 2017). The yeasts *Vishniacozyma victoriae* and *Filobasidium wieringae* (ex *Cryptococcus wieringae*) were also reported as moderate antagonists of several filamentous fungi (Hilber-Bodmer, Schmid, Ahrens, & Freimoser, 2017). Future research should investigate the interactions between these yeast species and grapevine foliar pathogens, including powdery mildew.

Contrary to our expectations, microbial networks were less sensitive to changes in agricultural practices than microbial communities, suggesting that NGB should not only consider network-level properties but also community-level properties. The α-properties of microbial networks (i.e. whole-network metrics; Pellissier et al., 2018) did not differ between cropping systems. Their variation was correlated with the network inference parameter, *P*, a methodological parameter corresponding to percentages of the most abundant taxa included in the network. Only two β-properties of microbial networks differed significantly between cropping systems, revealing a difference in microbial associations between organic and conventional vineyards. These differences remained significant when network pairwise comparisons were based on shared taxa only, suggesting that the differences between organic and conventional networks were due to re-associations of fungal taxa rather than a turnover of taxa. The functional redundancy of fungal taxa might account for the difference between networks. Taxonomically different communities, involving different interactions between members, may have similar functions (Louca et al., 2016) and protect plant health in a similar way. However, differences between networks might also be due to methodological hurdles that are discussed below and will have to be overcome in future NGB developments.

In our study, replicate microbial networks for the same cropping system had very few associations in common, revealing that the microbial networks we inferred from metabarcoding data lacked replicability. Three methodological parameters might account for this result. First, each network was built from 20 samples. Increasing the number of samples per network, to at least 25 (Berry & Widder, 2014b), might improve the replicability of microbial networks. Second, network nodes were amplicon sequence variants (ASVs) of the fungal ITS1 region, sequenced on an Illumina MiSeq platform. Despite their fine-scale taxonomic resolution (Callahan, McMurdie, & Holmes, 2017), ASVs may group together fungal strains with different interaction traits (McLaren & Callahan, 2018). A finer taxonomic resolution, achieved through the third-generation sequencing of the full ITS region (Nilsson et al., 2019), might improve the reliability of microbial networks (Kennedy, Cline, & Song, 2018). Third, our method of network reconstruction did not account for spatial variations in environmental conditions (i.e. microclimate or leaf traits) that could compromise inferences (Armitage & Jones, 2019). We did not measure environmental variations in our experimental system because the sampling design limited such variations. The vineyard plots were adjacent to each other and planted with grapevine clones. Moreover, we collected all leaves in less than two hours and controlled for the position of the sampled leaf on the vine. Our results suggest that this type of control might not be sufficient. Future developments of NGB approaches will have to include a higher number of samples per network, long-read sequencing and methods of network inference allowing the integration of environmental covariates (Chiquet et al., 2017, 2018; Cougoul et al., 2019; Ovaskainen et al., 2017; Tackmann et al., 2018).

Finally, our results show that consensus networks, built from several network replicates, can generate relevant hypotheses concerning microbial interactions. In our study, two negative associations were found to be common to all three replicates of the organic system. These negative associations were relevant according to current ecological knowledge. They involved two plant pathogens, *Neofusicoccum parvum* (commonly associated with grapevine trunk diseases; Bruez et al., 2014) and *Cladosporium ramotenellum* (associated with brown rot; Swett, Bourret, & Gubler, 2016), and two other species known to have antagonistic effects on plant pathogens, *Vishniacozyma victoriae* and *Aureobasidium sp.* (Lutz et al., 2013; Pertot et al., 2017). These results suggest that network inference is a promising tool to generate hypotheses that once tested will allow us to better understand disease regulation and perhaps discover biocontrol agent candidates (Poudel et al., 2016). Future research will however first have to determine whether it is better to build one big network with all samples or to merge several networks each based on a subset of the samples. In any case, confidence estimates of the links will have to be computed (Röttjers & Faust, 2018).

Despite these promising findings, this study indicates that the properties of NGS-based microbial networks cannot be used yet for monitoring the disease regulation services provided by the microbiota and that NGS-based microbial networks should be interpreted with caution (Carr, Diener, Baliga, & Gibbons, 2019). The replicated microbial association networks inferred from metabarcoding data were highly variable within each set of environmental conditions and generated only a few robust hypotheses concerning interactions between fungi, limiting for now their use to monitor the barrier effect of microbial interactions against pathogens. By contrast, community-level metrics revealed clear-cut changes in the plant microbiota in response to environmental change and reflected the disease status of the plant. Moreover, they were more statistically powerful than network-level metrics, because many samples were required to infer each microbial network replicate. Replicability of microbial networks is however likely to be improved in the near future, notably through developments in third-generation sequencing techniques and network inference methods, so microbial networks still hold promises for a reliable biomonitoring option (Karimi et al., 2017). Community-level data and network-level data, both analyzed using machine-learning approaches (e.g. Cordier et al., 2018), could even offer complementary insights into the ecosystem services provided by the plant microbiota.

## Acknowledgements

We thank Lucile Muneret, Andreas Makiola and all members of the ANR NGB Consortium (ANR-17-CE32-0011) for helpful scientific exchanges on next-generation biomonitoring and for their comments on the manuscript. We also thank Gregory Gambetta, Guilherme Martins, Frédéric Barraquand, Isabelle Lesur and Adrien Rush for useful comments on preliminary results. We thank the Genotoul sequencing facility (Get-PlaGe) for sequencing and the Genotoul bioinformatics facility (Bioinfo Genotoul) for providing computing and storage resources. We also thank Julie Sappa from Alex Edelman & Associates for English language revision of the first version of the manuscript. We thank the INRA MEM metaprogram (Meta-Omics of Microbial Ecosystems) for financial and scientific support. Sequencing was funded by the INRA MEM MetaBAR project and bioinformatic and statistical analyses were performed as part of the INRA MEM Learn-biocontrol project. Additional funding was received from the LABEX COTE (ANR-10-LABX-45), the LABEX CEBA (ANR-10-LABX-25-01) and INRA EcoServ metaprogram on ecosystem services (IBISC project) and the Aquitaine Region (Athene project, n°2016-1R20301-00007218). CP’s PhD grant was funded by the INRA and Bordeaux Sciences Agro (BSA). The management of the experimental site was partly funded by the AFB (French Agency for Biodiversity) within the DEPHY network.

## Data Accessibility Statement

The raw sequence data were deposited in Dataverse and are available in the FASTQ format at https://doi.org/10.15454/3DPFNJ while the filtered ASV table is available at https://doi.org/10.15454/WOICSE. The code is available as an archive at https://doi.org/10.15454/NSHUAQ.

## Author Contributions

CP performed the bioinformatic and statistical analyses and wrote the first draft of the manuscript. JV and CV performed the sampling and processed the leaf samples. LD managed the sampling site and provided data on phytosanitary treatments and disease symptoms. JV performed the DNA extractions and amplifications. MB coordinated the creation of the fungal mock community and the sequencing of all samples. DB contributed to the original idea of the study and data interpretation. CV designed the study, supervised the analyses and made a major contribution to the writing of the final draft. All authors discussed the preliminary version of the results and revised the manuscript.

## Supplementary materials

**Figure S1.**
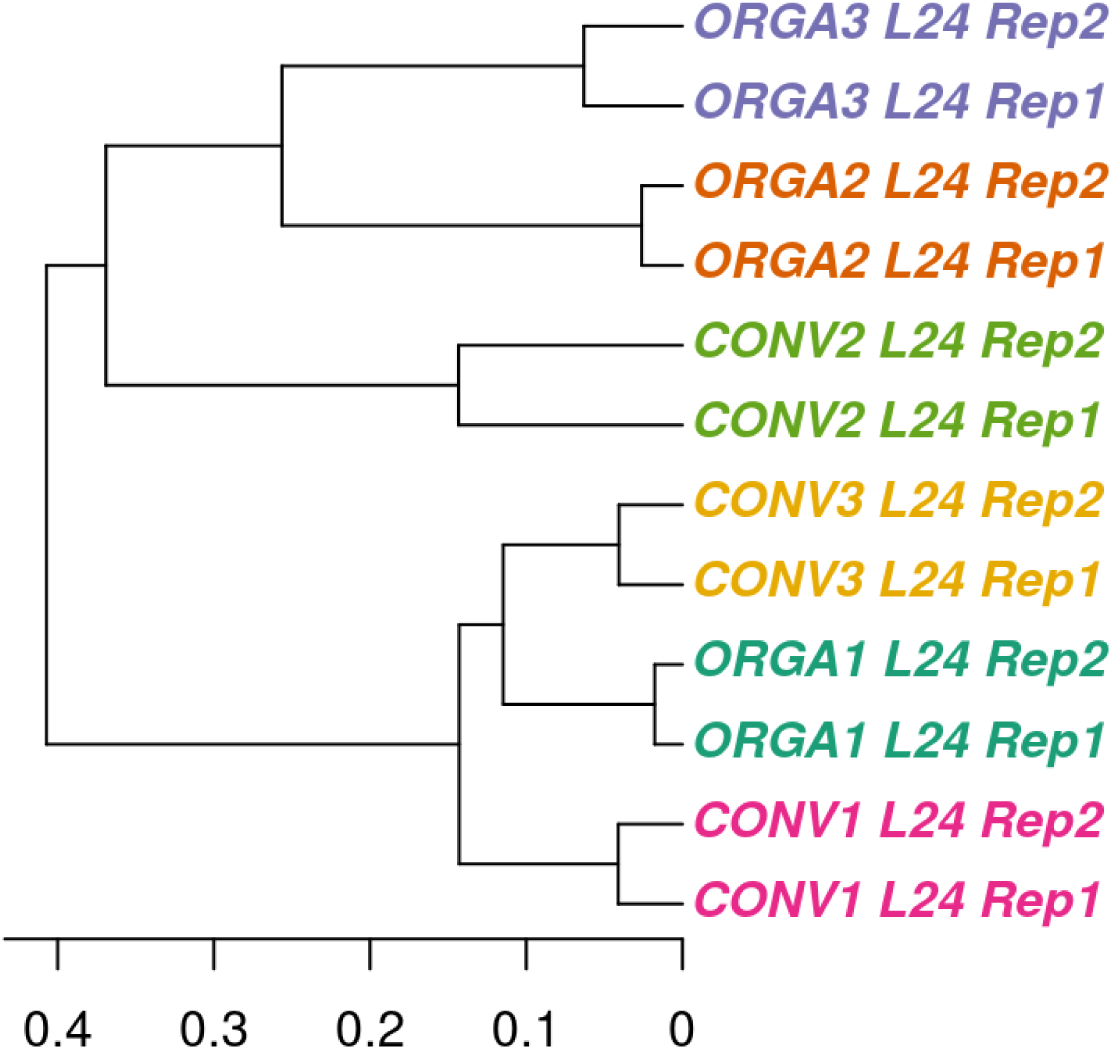
Dendrogram plot of compositional dissimilarities between technical replicates for sequencing. Technical replicates were created by splitting six lots of PCR products in half and sequencing the two halves independently. The PCR products used were those corresponding to leaf 24 (L24) of the six plots studied (ORGA1, ORGA2, ORGA3, CONV1, CONV2, CONV3; see Figure 5). Compositional dissimilarities between samples were computed with the binary Jaccard index. The dendrogram was built using a hierarchical clustering algorithm (complete linkage method). Compositional dissimilarities between the two technical replicates of the same sample were significantly smaller than the dissimilarities among samples (PERMANOVA: *F* = 39.98; R2 = 0.97; *p* = 0.001).

**Figure S2.**
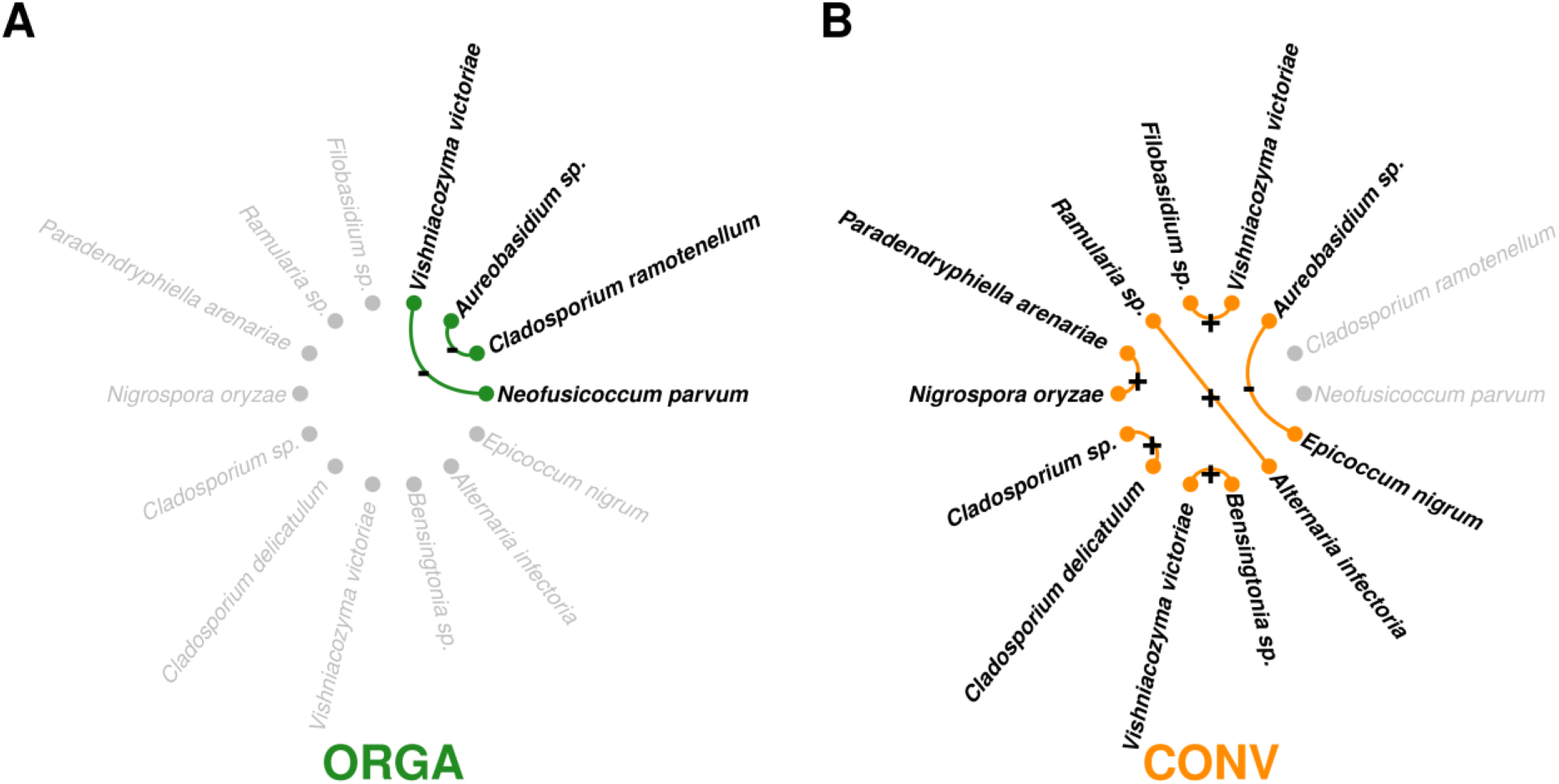
Consensus fungal networks for (A) the organic (ORGA) and (B) the conventional (CONV) cropping systems. Network nodes represent fungal ASVs and links represent significant positive (+) or negative (-) associations common to the three network replicates (Fig. 6A and 6B, respectively). The fungal ASVs absent from a network are indicated in gray.

**Table S1.**
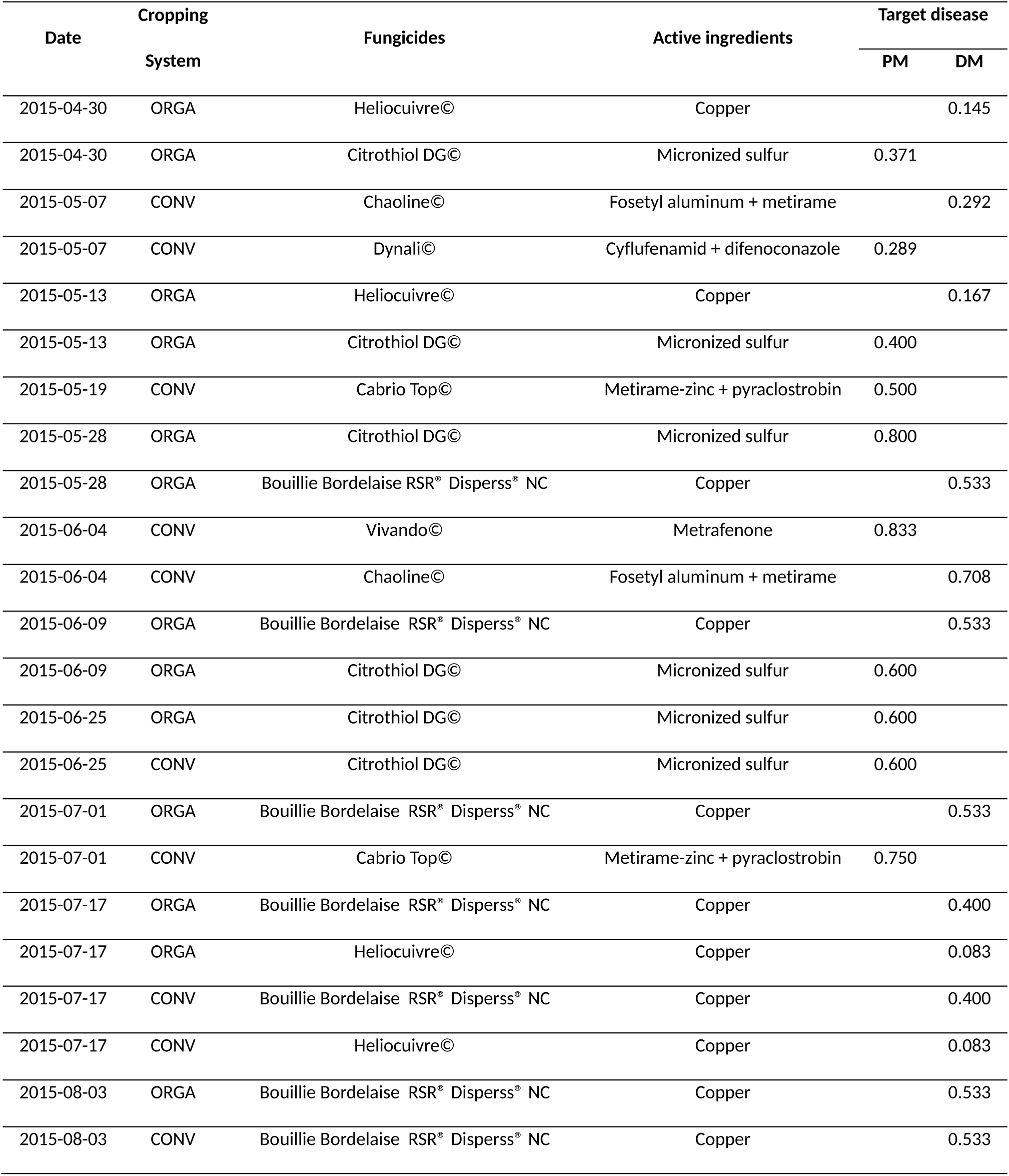
List of phytosanitary products and active ingredients applied in the year of the sampling campaign, together with the normalized dose or the treatment frequency index. PM = powdery mildew (caused by the fungal pathogen *Erysiphe necator*) and DM = downy mildew (caused by the oomycete pathogen *Plasmopara viticola*). Leaf sampling was performed on September 10 2015 (more than one month after the last phytosanitary treatment and a couple of hours before grape harvest). The treatment frequency index did not differ between cropping systems (ANOVA: df = 21; *F* = 0.436; *p* = 0.516).

**Table S2.** Effect of cropping system —conventional (CONV) *versus* organic (ORGA) — on the incidence and severity of foliar disease symptoms at harvest time (2015-09-07). Disease incidence is defined as the percentage of leaves displaying symptoms, whereas disease severity is defined as the percentage leaf damage. Symptom incidence and severity were estimated visually on 40 grapevines for each plot (40 × 3 per cropping system). The mean values are reported for each cropping system as a percentage. Wald χ^2^ tests were used for comparisons after linear mixed model analysis with cropping system as a fixed effect and block as a random effect.

| Disease | | ORGA (%) | CONV (%) | $\chi^2$ | p-value |
| --- | --- | --- | --- | --- | --- |
| Downy | Incidence | 0.749 | 0.688 | 0.57 | 0.450 |
|  | Severity | 0.037 | 0.030 | 1.93 | 0.164 |
| Powdery | Incidence | 0.113 | 1.346 | 12.49 | **<0.001** |
|  | Severity | 0.003 | 0.102 | 7.97 | **0.005** |
| Black rot | Incidence | 0.188 | 0.354 | 19.02 | **<0.001** |
|  | Severity | 0.007 | 0.014 | 5.49 | **0.019** |

**Table S3.**
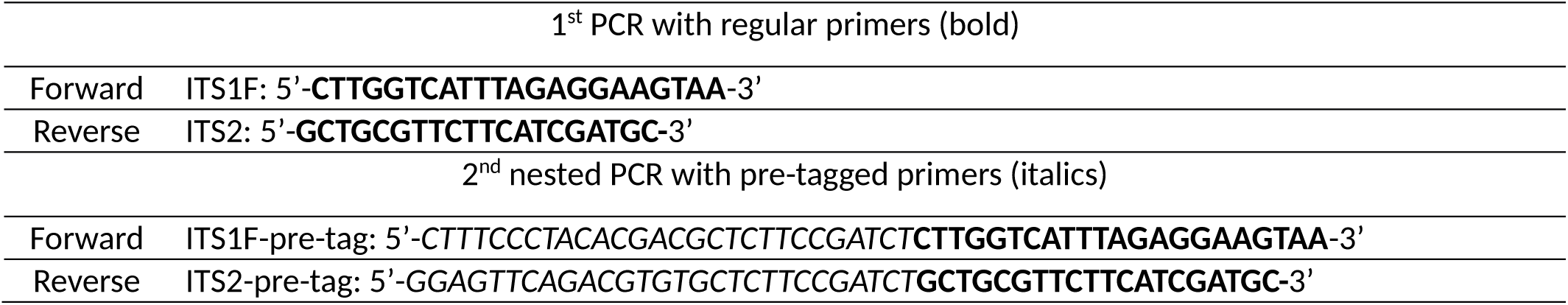
Primer pairs used to amplify the fungal ITS1 region.

**Table S4.** List of network α-properties calculated in this study.

| **Network metric** | **Definition** | **Reference** |
| --- | --- | --- |
| Number of links (L) | Total number of links |  |
| Connectance (C) | Fraction of the total number of possible links actually realized | Coleman and Moré 1983<br>10.1137/0720013 |
| Number of connected components (CC) | Number of groups of nodes connected together | Martinez 1992<br>10.1086/285382 |
| Diameter (DIA) | The longest of all the shortest paths between two nodes | Barabási et al., 2000<br>10.1016/S0378-4371(00)00018-2 |
| Mean node degree (DEG) | Mean number of links per node | Martinez 1992<br>10.1086/285382 |
| Proportion of negative links (NLR) | Proportion of links for which the SparCC correlation is negative | Faust et al., 2015<br>10.3389/fmicb.2015.01200 |

**Table S5.** Effect of cropping system on the α-properties of fungal association networks. Properties (as defined in Table S4) were compared between cropping systems for every value of the percentage *P* of the most abundant ASVs used for network inference. The *U* and *p*-values of Wilcoxon rank-sum tests are reported. The *p*-value is not available (NA) for situations in which property values were equal for all networks. The *p*-values after Benjamini-Hochberg adjustment are not reported because all were equal to one.

| $P$ (%) | L | CC | DIA | C | DEG | NLR |
| --- | --- | --- | --- | --- | --- | --- |
| 10 | $U = 3; p = 0.658$ | $U = 5; p = 1$ | $U = 6; p = 0.505$ | $U = 3; p = 0.663$ | $U = 2; p = 0.383$ | $U = 5; p = 1$ |
| 20 | $U = 1; p = 0.19$ | $U = 6.5; p = 0.48$ | $U = 8; p = 0.157$ | $U = 2; p = 0.383$ | $U = 1; p = 0.19$ | $U = 9; p = 0.081$ |
| 30 | $U = 1; p = 0.19$ | $U = 4.5; p = \text{NA}$ | $U = 6; p = 0.619$ | $U = 1; p = 0.19$ | $U = 1; p = 0.19$ | $U = 6; p = 0.663$ |
| 40 | $U = 1; p = 0.19$ | $U = 4.5; p = \text{NA}$ | $U = 7; p = 0.302$ | $U = 2; p = 0.383$ | $U = 1; p = 0.19$ | $U = 5; p = 1$ |
| 50 | $U = 3; p = 0.663$ | $U = 4.5; p = \text{NA}$ | $U = 7; p = 0.302$ | $U = 3; p = 0.663$ | $U = 3; p = 0.663$ | $U = 6; p = 0.663$ |
| 60 | $U = 2; p = 0.383$ | $U = 4.5; p = \text{NA}$ | $U = 4.5; p = 1$ | $U = 2; p = 0.383$ | $U = 1; p = 0.19$ | $U = 5; p = 1$ |
| 70 | $U = 3; p = 0.663$ | $U = 4.5; p = \text{NA}$ | $U = 9; p = 0.047$ | $U = 2; p = 0.383$ | $U = 2; p = 0.383$ | $U = 5; p = 1$ |
| 80 | $U = 3; p = 0.663$ | $U = 4.5; p = \text{NA}$ | $U = 6; p = 0.505$ | $U = 4; p = 1$ | $U = 3; p = 0.663$ | $U = 4; p = 1$ |
| 90 | $U = 4; p = 1$ | $U = 4.5; p = \text{NA}$ | $U = 6; p = 0.505$ | $U = 4; p = 1$ | $U = 3; p = 0.663$ | $U = 3; p = 0.663$ |
| 100 | $U = 3; p = 0.663$ | $U = 4.5; p = \text{NA}$ | $U = 4.5; p = \text{NA}$ | $U = 2; p = 0.383$ | $U = 4; p = 1$ | $U = 4; p = 1$ |

**Table S6.** Effect of the percentage *P* of the most abundant ASVs used for network inference on the α-properties of fungal association networks. Spearman’s correlation coefficient and the results of Spearman’s rank correlation tests are reported for each network property. The *p*-values are reported after Benjamini-Hochberg adjustment.

| Property | Correlation ($\rho$) | S | *p*-value |
| --- | --- | --- | --- |
| L | 0.98 | 862 | <0.001 |
| CC | -0.61 | 57987 | <0.001 |
| DIA | -0.85 | 66635 | <0.001 |
| C | -0.67 | 60144 | <0.001 |
| DEG | 0.95 | 1635 | <0.001 |
| NLR | -0.56 | 56116 | <0.001 |

